# Choosing what we like vs liking what we choose: How choice-induced preference change might actually be instrumental to decision-making

**DOI:** 10.1101/661116

**Authors:** Douglas Lee, Jean Daunizeau

## Abstract

For more than 60 years, it has been known that people report higher (lower) subjective values for items after having selected (rejected) them during a choice task. This phenomenon is coined “choice-induced preference change” or CIPC, and its established interpretation is that of “cognitive dissonance” theory. In brief, if people feel uneasy about their choice, they later convince themselves, albeit not always consciously, that the chosen (rejected) item was actually better (worse) than they had originally estimated. While this might make sense from a pragmatic psychological standpoint, it is challenging from a theoretical evolutionary perspective. This is because such a cognitive mechanism might yield irrational biases, whose adaptive fitness would be unclear. In this work, we assume that CIPC is mostly driven by the refinement of option value representations that occurs *during* (and not *after*) difficult choices. This makes CIPC the epiphenomenal outcome of a cognitive process that is instrumental to the decision. Critically, our hypothesis implies novel predictions about how observed CIPC should relate to two specific meta-cognitive processes, namely: choice confidence and subjective certainty regarding pre-choice value judgments. We test these predictions in a behavioral experiment where participants rate the subjective value of food items both before and after choosing between equally valued items; we augment this traditional design with reports of choice confidence and subjective certainty about value judgments. The results confirm our predictions and provide evidence against the standard post-choice cognitive dissonance reduction explanation. We then discuss the relevance of our work in the context of the existing debate regarding the putative cognitive mechanisms underlying cognitive dissonance reduction.

## INTRODUCTION

The causal relationship between choices and subjective values goes both ways. By definition, choices are overt expressions of subjective values, which is the basis of decision theory (Slovic, Fischhoff & Lichtenstein, 1977). However, one’s choices also influence one’s values, such that actions or items seem to acquire value simply because one has chosen them. Such “choice-induced preference change” (CIPC) has been repeatedly demonstrated via the so-called “free-choice paradigm” (Brehm, 1956). Here, people rate the pleasantness of (e.g., food) items both before and after choosing between pairs of equally pleasant items. Results show that the post-choice pleasantness ratings of chosen (rejected) options are typically higher (lower) than their pre-choice pleasantness ratings, which has traditionally been taken as empirical evidence for the existence of a “cognitive dissonance” reduction mechanism, triggered by difficult choices (see Harmon-Jones & Harmon-Jones, 2007; Izuma & Murayama, 2013 for reviews). An established variant of this interpretation is that people rationalize their choice *ex post facto* as they think along the lines of, “I chose (rejected) this item, so I must have liked it better (worse) than the other one,” and hence adjust their internal values accordingly (Bem, 1967). Over the past decade, neuroimaging studies have demonstrated that the act of choosing between similarly-valued options causes changes in the brain’s encoding of subjective values (Izuma et al, 2010; Voigt et al, 2018). This has lent neurobiological support to the theory, and cognitive dissonance reduction is now the popular explanation behind a broad variety of important irrational sociopsychological phenomena, ranging from, for example, post-vote political opinion changes (Beasley & Joslyn, 2001) to post-violence hostile attitude worsening (Acharya, Blackwell & Sen, 2015).

This is not to say, however, that the theory of cognitive dissonance reduction has remained unchallenged. The first issue is theoretical in essence. In brief, it is unclear why evolutionary pressure would have favored post-choice cognitive dissonance reduction mechanisms, given that they could eventually induce irrational cognitive biases that have no apparent adaptive fitness (Akerlof and Dickens, 1982; Gilad et al., 1987; Perlovsky, 2013). For example, in the context of evidence-based decision making, standard cognitive dissonance theory predicts the appearance of confirmation and overconfidence biases. This is simply because weak beliefs should be reinforced by subsequent choices, despite the lack of any additional piece of evidence (Allahverdyan and Galstyan, 2014; Navajas et al., 2016). At least in principle, cognitive dissonance reduction may of course have other behavioral consequences that would overcompensate for the adverse selective pressure on cognitive biases. Now if it possessed such adaptive fitness, then it would undoubtedly be expressed in many other animal species. This is, however, an unresolved issue in the existing ethological literature (Egan, Santos & Bloom, 2007; West et al, 2010). Second, the main experimental demonstration of cognitive dissonance has also been challenged on statistical grounds. In 2010, Chen and Risen reported a methodological issue in the way CIPC had typically been measured and explained. The basic idea was that simple random variability in repeated value ratings could confound classical measures of CIPC in the context of the free-choice paradigm. The authors provided a detailed mathematical explanation for how such a statistical confound might eventually cause an apparent CIPC (see Chen & Risen, 2010 for details), and introduced a clever control condition. Here, both first and second value ratings are provided before any choice is ever made, thus precluding choice from causally influencing reported subjective values. Results show that significant CIPC occurs regardless of whether the choice is made before or after the second rating. Although this supports the validity of the authors’ statistical criticism, subsequent studies also demonstrated that the magnitude of CIPC is significantly greater when the choice is made before the second value rating (Salti et al, 2014; Coppin et al, 2013; Coppin et al, 2012; Sharot et al, 2012). Taken together, the current theoretical and empirical bases of CIPC do not yet provide a straightforward portrait of why and how choice may influence subjective values.

Interestingly, recent neuroimaging evidence suggests that, in the context of typical free-choice paradigms, preference changes occur *during* the decision, not *after* it (Jarcho, Berkman, & Lieberman, 2010; Colosio et al, 2017). Recall that people are reluctant to make a choice that they are not confident about (De Martino et al, 2013). But envisioning competing possibilities during a choice provides a new context that highlights the unique aspects of the alternative options (Tversky & Thaler, 1990; Lichtenstein & Slovic, 2006; Warren, McGraw, & Van Boven, 2011). In turn, the act of choosing may change preferences by reappraising aspects of choice options that may not have been considered thoroughly before (Sharot et al., 2009). In other words, preferences may change during deliberation in order to facilitate the choice process (Simon, Stenstrom & Read, 2015; Simon, Krawczyk & Holyoak, 2004; Simon & Holyoak, 2002). This is important, because it allows for the possibility that preference changes may be instrumental for the process of decision making, which would resolve most theoretical concerns. This is the essence of our working hypothesis. We reason that decision difficulty drives people to reassess the values of the alternative options before committing to a particular choice. The ensuing refinement of internal value representations will eventually raise choice confidence enough to trigger the decision, which may or may not be aligned with pre-choice value ratings. Critically for our theory, the more difficult the decision, the more deliberation and potential reassessment of value representations, the more likely a change of mind and the related CIPC. This would make CIPC the epiphenomenal outcome of a cognitive process that is instrumental to the decision.

Importantly, our working hypothesis makes two original predictions that deviate from standard post-choice cognitive dissonance reduction theory. Recall that the magnitude of CIPC is known to increase with the absolute difference between pre-choice option values, which is typically taken as a proxy for choice difficulty (Izuma and Murayama, 2013; Sharot et al., 2012). We argue, however, that choice difficulty is better defined in terms of the similarity of value representations. The critical difference is that value representations may be uncertain, i.e. the feeling of liking and/or wanting a given choice option may be imprecise. In other terms, subjective estimates of choice difficulty derive from both value difference and metacognitive judgments about value uncertainty. Our first prediction regards the impact of the latter. In particular, we predict that CIPC should increase with subjective uncertainty regarding pre-choice value judgment. This is because pre-choice value uncertainty raises choice difficulty, which calls for more value reassessment. In contrast, standard post-choice cognitive dissonance reduction theory predicts that post-choice dissonance should be highest when pre-choice values were *a priori* certainly equal, i.e. for choices with minimal value distance and maximal value certainty. Second, we predict that CIPC should positively correlate with choice confidence, controlling for the impact of decision difficulty. This is because, under our hypothesis, CIPC indirectly signals a successful improvement of choice confidence, due to reassessed values spreading apart. In contrast, standard dissonance reduction theory would posit that choices made with low confidence should trigger the strong aversive dissonance feelings that eventually lead to CIPC. Note that we can rule out variants of Chen and Risen’s statistical confound by showing that, if anything, evidence for both predictions is weaker when the choice is made after the second value rating.

## METHODS

We adapted the experimental design of Chen and Risen (2010), which includes two groups of participants. The so-called RCR (Rating, Choice, Rating) group of participants was asked to rate the value of a series of items both before and after making paired choices. In contrast, the RRC (Rating, Rating, Choice) group of participants rated the items twice before making the paired choices. As we will see, comparisons between the RCR and the RRC (control) group serve to rule out variants of Chen and Risen’s statistical confound. In our adapted experimental design, when evaluating subjective values, participants now also rated their subjective certainty regarding their value judgment. In addition, they also reported how confident they were when making paired choices. This allows us to assess the impact of both subjective uncertainty about value judgments and choice confidence on CIPC.

### Participants

A total of 123 people participated in this study. The RCR group included 65 people (45 female; age: mean=29, stdev=9, min=19, max=53). The RRC group included 58 people (34 female; age: mean=33, stdev=11, min=18, max=55). All participants were native French speakers. Each participant was paid a flat rate of 12€ as compensation for one hour of time.

### Materials

We wrote our experiment in Matlab, using the Psychophysics Toolbox extensions (Brainard, 1997). The experimental stimuli consisted of 108 digital images, each representing a distinct sweet snack item (including cookies, candies, and chocolates). Prior to the experiment, participants received written instructions about the sequence of tasks, including typical visual examples of rating and choice trials.

### Experimental Design

The experiment was divided into three sections, following the classic Free-Choice Paradigm protocol: Rating #1, Choice, Rating #2 (RCR group) or Rating #1, Rating #2, Choice (RRC group). Note that only in the RCR group do Rating #1 and Rating #2 correspond to pre-choice and post-choice ratings. Participants underwent a brief training session prior to the main testing phase of the experiment. There was no time limit for the overall experiment, nor for the different sections, nor for the individual trials. Within-trial event sequences are described below (Figure 1).

**Figure 1:**
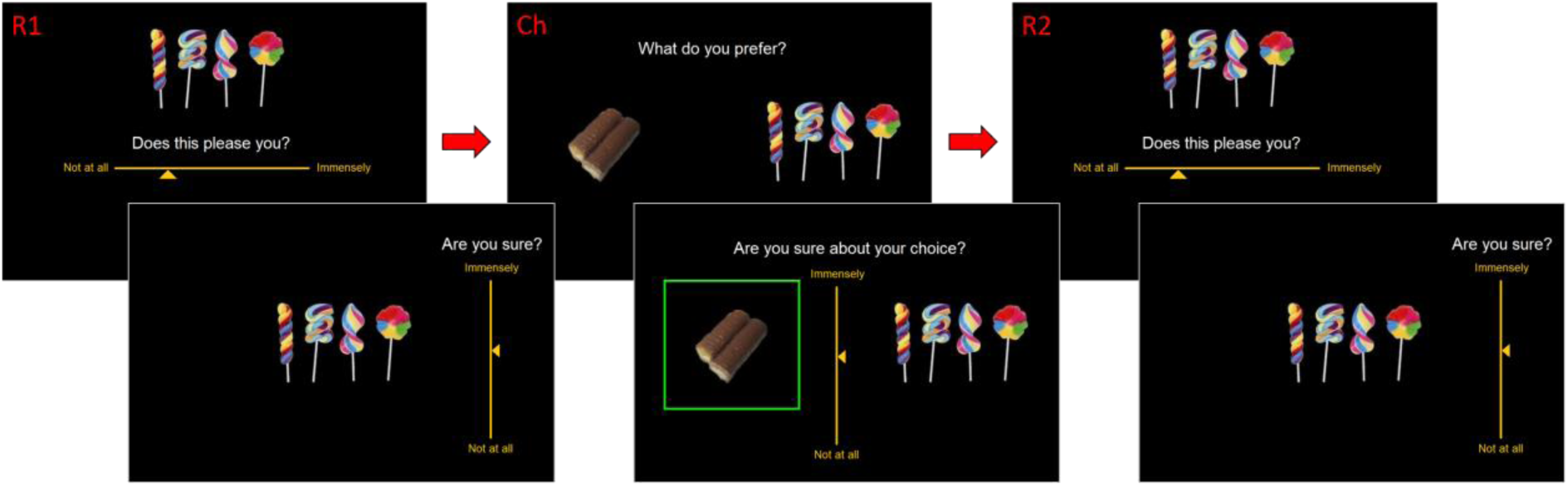
Experiment Tasks: Rating #1, Choice, Rating #2.

#### Rating

Participants rated the stimulus items in terms of how much each item pleased them. The entire set of stimuli was presented to each participant, one at a time, in a random sequence (randomized across participants). At the onset of each trial, a fixation cross appeared at the center of the screen for 750ms. Next, a solitary image of a food item appeared at the center of the screen. Participants had to respond to the question, “Does this please you?” using a horizontal Likert scale (from “not at all” to “immensely”) to report their subjective valuation of the item. Participants then had to respond to the question, “Are you sure?” using a vertical Likert scale (from “not at all” to “immensely”) to indicate their level of subjective uncertainty regarding the preceding value judgment. At that time, the next trial began.

#### Choice

Participants chose between pairs of items in terms of which item they preferred. The entire set of stimuli was presented to each participant, one pair at a time, in a random sequence of pairs. Each item appeared in only one pair. The algorithm that created the choice pairs first sorted all items into 10 bins, then paired off (at least) half of the items within each bin, then paired off all remaining items across bins. This ensured that at least half of choices would be between items of similar subjective value (value rating difference < 1/10 of the full rating scale, as shown in previous studies to cause CIPC), but that a substantial portion would be associated with greater value differences.

At the onset of each trial, a fixation cross appeared at the center of the screen for 750ms. Next, two images of snack items appeared on the screen: one towards the left and one towards the right. Participants had to respond to the question, “What do you prefer?” using the left or right arrow key. Participants then had to respond to the question, “Are you sure about your choice?” using a vertical Likert scale to report their level of confidence in the preceding choice. At that time, the next trial began.

## RESULTS

Before testing our hypothesis (against both statistical confounds and standard post-choice cognitive dissonance reduction theory), we performed a number of simple data quality checks. First, we assessed the test-retest reliability of both value judgments and their associated certainty reports. For each participant, we thus measured the correlation between ratings #1 and #2 (across items). We found that both ratings were significantly reproducible (value ratings: correlation = 0.862, 95% CI [0.838, 0.886], p<0.001; certainty ratings: correlation = 0.472, 95% CI [0.409, 0.535], p<0.001). Second, we asked whether choices were consistent with value ratings #1. For each participant, we thus performed a logistic regression of paired choices against the difference in value ratings. We found that the balanced prediction accuracy was beyond chance level (group mean = 0.685, 95% CI [0.666, 0.703], p<0.001). Third, we checked that choice confidence increases both with the absolute value difference between paired items, and with the mean certainty reports about value judgments (of the paired items). For each participant, we thus performed a multiple linear regression of choice confidence against absolute value difference and mean judgment certainty (ratings #1). A random effect analysis shows that both have a significant effect at the group level (R^2^ = 0.223, 95% CI [0.188, 0.258]; absolute value difference: GLM beta = 9.046, 95% CI [7.841, ∞], p<0.001; mean judgment certainty: GLM beta = 3.157, 95% CI [2.137, ∞], p<0.001). Fourth, we asked whether we could reproduce previous findings that CIPC is higher in the RCR group than in the RRC group. For each participant, we thus measured the magnitude of CIPC in terms of the so-called “spreading of alternatives” (SoA), calculated as the mean difference in value rating gains between chosen and unchosen items (SoA = [rating#2-rating#1]_chosen_ - [rating#2-rating#1]_unchosen_). As expected, we found that SoA is significant in both groups (RCR group: SoA = 5.033, 95% CI [4.118, 5.949], p<0.001; RRC group: SoA = 2.635, 95% CI [2.047, 3.224], p<0.001). In addition, SoA is significantly higher in the RCR group than in the RRC group (SoA difference = 2.398, p<0.001) (Figure 2).

**Figure 2:**
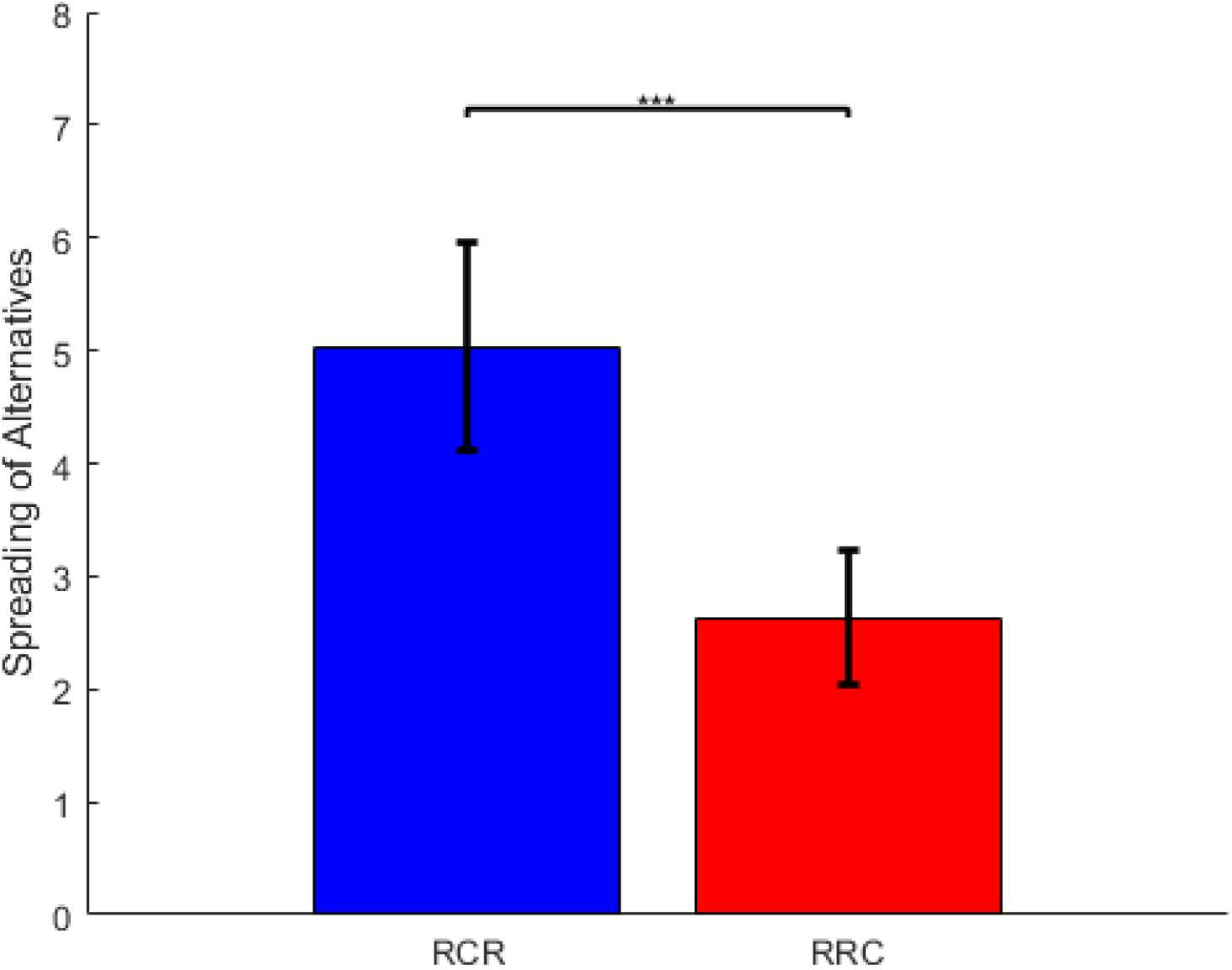
Mean and 95% CI for spreading of alternatives in the RCR and RRC groups.

In what follows, and unless stated otherwise, we focus on the RCR group of participants. Recall that, under our hypothesis, the deliberation that takes place during the decision process is expected to cause a refinement of internal value representations until a target level of choice confidence is met and the decider commits to a choice. To begin with, we thus asked whether certainty about value judgments improved after the choice had been made. For each participant, we thus estimated the mean difference between post-choice and pre-choice certainty reports (across all items). A random effect analysis then shows that post-choice certainty reports are significantly higher than pre-choice certainty reports (certainty increase = 3.781, 95% CI [1.810, 5.752], p<0.001). This finding supports our claim but does not provide evidence for or against classical post-choice cognitive dissonance reduction theory. We then asked whether post-choice ratings better predict choice (and choice confidence) than pre-choice ratings. First, we performed another logistic regression of paired choices, this time against the difference in post-choice value ratings (ratings #2). The ensuing choice prediction accuracy is higher than with pre-choice value ratings (accuracy = 0.787, 95% CI [0.770, 0.804], accuracy gain = 0.103, 95% CI [0.082, 0.124], p<0.001) (Figure 3). Second, we regressed choice confidence, this time against post-choice absolute value difference and mean judgment certainty. The ensuing amount of explained variance in choice confidence reports is higher than with pre-choice ratings (R^2^ = 0.245, 95% CI [0.209, 0.281], R^2^ gain = 0.022, 95% CI [0.001, 0.042], p=0.02) (Figure 3). When testing for the significance of differences in pre-choice and post-choice regression parameters, we found that this gain in explanatory power is more likely to be due to value ratings (GLM beta difference = 0.763, 95% CI [0.180, ∞], p=0.018) than to certainty reports (GLM beta difference = 0.158, 95% CI [-0.451, ∞], p=0.34). These results are important, because they validate basic requirements of our pre-choice CIPC hypothesis. However, they are equally compatible with both pre-choice and post-choice CIPC mechanisms. Next, we focus on testing the two specific predictions regarding the relationship between CIPC and meta-cognitive processes, which discriminate mechanisms of pre-choice value refinements from post-choice cognitive dissonance reduction.

**Figure 3:**
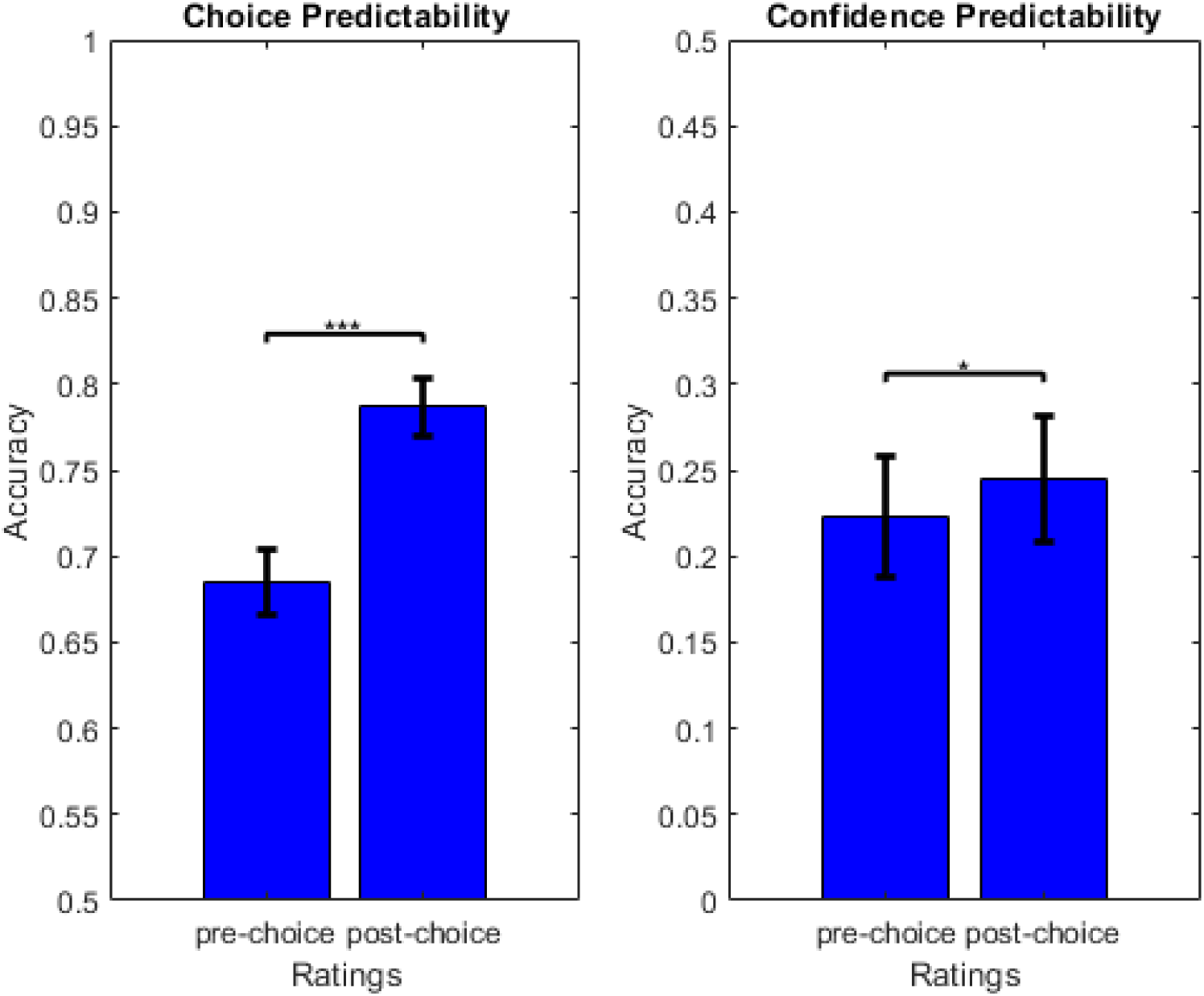
Mean and 95% CI for choice predictability (left) and choice confidence predictability (right), based on pre-vs post-choice ratings.

We now assess the statistical relationships between CIPC and both choice difficulty and choice confidence. For each participant, we performed a multiple linear regression of SoA onto absolute difference in pre-choice value ratings, mean pre-choice judgment certainty reports, and choice confidence (Figure 4). As expected, a random effect analysis on the ensuing parameter estimates shows that SoA significantly decreases with the absolute difference in pre-choice value ratings (GLM beta = −0.295, 95% CI [-∞, −0.265], p<0.001). More importantly, we found that SoA significantly decreases with pre-choice judgment certainty (GLM beta = - 0.065, 95% CI [-∞, −0.036], p<0.001) and increases with choice confidence (GLM beta = 0.200, 95% CI [0.170, ∞], p<0.001). The latter findings support our hypothesis, and are incompatible with classical post-choice cognitive dissonance reduction theory.

**Figure 4:**
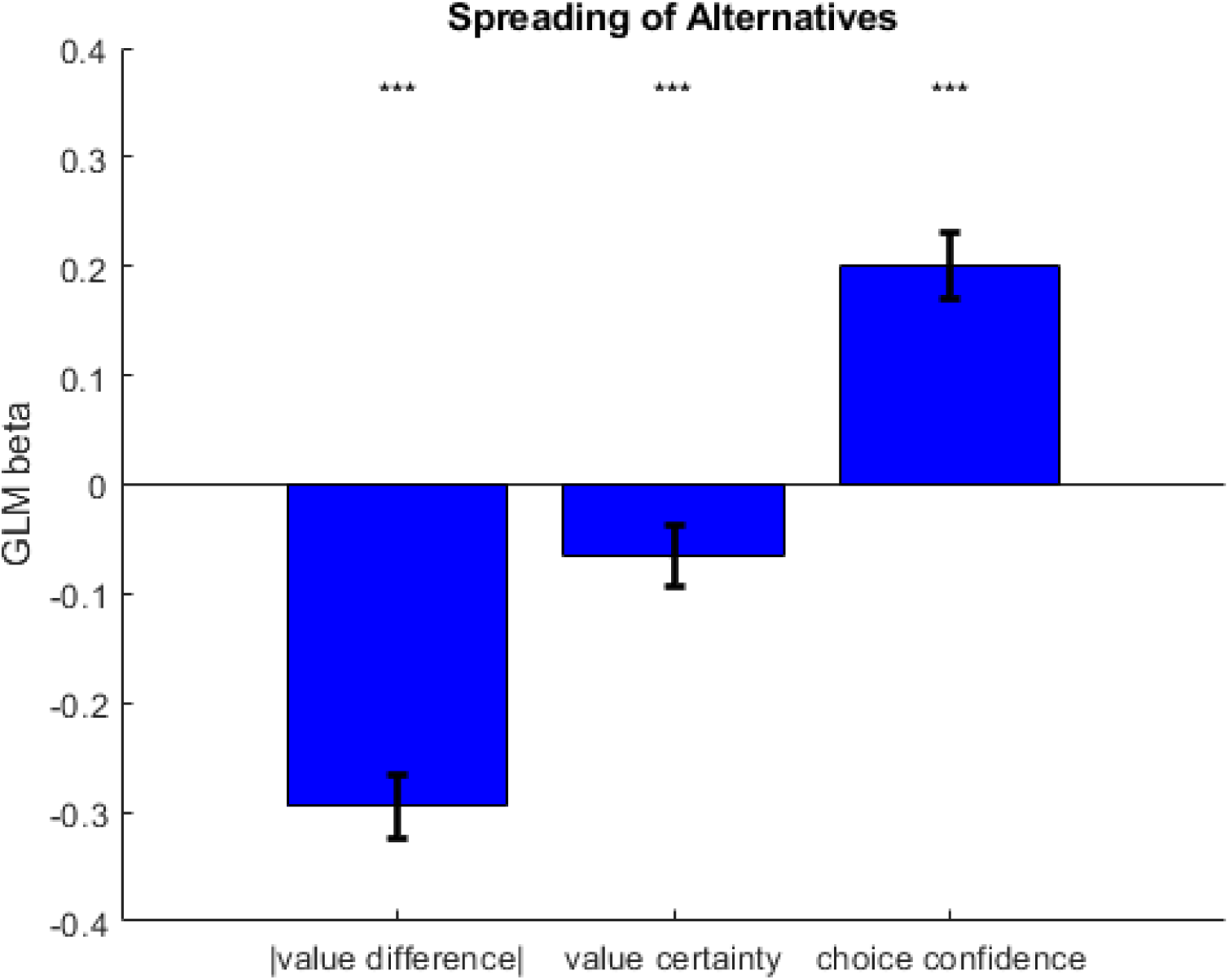
Mean and 95% CI for GLM beta weights of absolute value difference, average value certainty (within a choice pair), choice confidence on spreading of alternatives.

Finally, we aimed at ruling out statistical confounds. This can be done by showing that if the above statistical relationships exist in the RRC group, they should be significantly weaker than in the RCR group. We thus performed the above analyses on data acquired in participants from the RRC group, which we compared to the RCR group of participants using standard random effect analyses. First, the gain in choice prediction accuracy (from rating #1 to rating #2) is significantly higher in the RCR group than in the RRC group (accuracy gain difference = 0.0362, p=0.008). Second, and most importantly, both the impact of absolute pre-choice value difference (GLM beta diff = −0.095, p=0.0039) and choice confidence (GLM beta diff = 0.044, p=0.049) on SoA are significantly higher in the RCR than in the RRC group. Note that some comparisons between the two groups turned out not to be significant (gain in subjective certainty regarding value judgments: p=0.17, gain in confidence prediction accuracy: p=0.58, impact of subjective value certainty on SoA: p=0.096). Nevertheless, taken together, these findings are unlikely under a chance model of random variations in value ratings.

## DISCUSSION

In this work, we have presented empirical evidence that challenges standard interpretations of CIPC, in particular: post-choice cognitive dissonance reduction theory (and its self-perception variants). However, we would like to highlight that we do not dispute the concept of cognitive dissonance itself, nor even the idea that cognitive dissonance is the root cause of CIPC. In fact, our view only differs from standard post-choice cognitive dissonance reduction theory in one single aspect, namely: the temporal dynamics of the dissonance reduction. According to standard post-choice cognitive dissonance reduction theory, choices made with low confidence trigger strong aversive dissonance feelings that are resolved by retrospectively matching internal value representations to the choice. We would rather say that no choice commitment is made until internal value representation refinements allow choice confidence to reach a satisfying (non-aversive) level. This idea is a simple extension of the so-called Action-based model of cognitive dissonance (Harmon-Jones, Harmon-Jones, & Levy, 2015; Harmon-Jones, Amodio, & Harmon-Jones, 2009), and of recent attempts to formalize cognitive dissonance in terms of neurocognitive theories of predictive processing in the action-perception loop (Kaaronen, 2018). That cognitive dissonance reduction occurs during the decision process (and not after) is critical, however, because it is now endowed with a clear functional purpose (namely: improving decision accuracy).

Note that one may challenge our interpretation of the observed relationship between confidence and CIPC, in terms of evidence against post-choice CIPC. For example, one may argue that CIPC might occur after people commit to a choice, but before they get a feel for how confident they are about that choice. This sounds paradoxical however, in the sense that experiencing cognitive dissonance in this context simply means feeling uneasy about one’s choice, i.e. lacking confidence about it. In any case, this line of reasoning cannot apply to the observed impact of value certainty on CIPC. Recall that we probe metacognitive judgments about value certainty before the choice, using rating scales at the time when each item is presented (immediately after first-order value judgments). Therefore, the relationship between CIPC and value certainty that we disclose empirically cannot derive from metacognitive processes that occur after the choice has been made.

Our results apparently contradict the recent finding that CIPC only occurs for choices where the agent is later able to recall which option was chosen and which was rejected (Salti et al, 2014). This is because remembering choices has no causal role under our pre-choice value refinement hypothesis. This contrasts with standard cognitive dissonance reduction theory, where post-choice option re-evaluation requires the memory trace of relevant choices. This interpretation is compatible with the observation that, when re-evaluating items after the choice, activity in the hippocampus discriminates between remembered and non-remembered choices (Chammat et al, 2017). These two studies thus provide apparent evidence that CIPC occurs after the choice has been made. However, we contend that this theory remains unsupported until empirical evidence is found for memory traces of information that is critical for post-choice CIPC (namely: whether an option was chosen or rejected, what was the option’s pre-choice value, and which option was the alternative during the relevant choice). In addition, the relationship between CIPC and memory might be confounded by choice difficulty. In brief, the more difficult a decision is, the more value reassessment it will eventually trigger, the more likely the agent is to remember his/her choice. Alternatively, post-choice reports of internal values may rely on slightly unstable episodic memory traces of intra-choice CIPC. The latter scenario is actually compatible with the fact that activity in the left dorsolateral prefrontal cortex (during choice) predicts the magnitude of CIPC only when the choices are remembered (Voigt et al, 2018), and also with the intra-choice CIPC interpretation of the causal impact of post-choice activity perturbations (see below). In any case, either or both of these scenarios would explain why intra-choice CIPC might exhibit an apparent (non-causal) relationship with choice memory. Finally, note that the causal implication of memory is inconsistent with the assessment of amnesic patients, who exhibit normal CIPC despite severe deficits in choice memory (Liebermann et al, 2001).

Even though the cognitive architecture that underlies intra-choice cognitive dissonance reduction is yet to be disclosed, recent neuroimaging findings shed light on the question of whether CIPC occurs during or after the decision. On the one hand, a few brain stimulation studies suggest that perturbing brain activity *after the choice* (in particular: in the left dorsolateral and/or posterior medial frontal cortices) disrupts the observed CIPC (Mengarelli et al, 2013; Izuma et al, 2015). Although compatible with post-choice CIPC, such causal effects can be due to the post-choice disturbance of value representations that resulted from intra-choice CIPC. On the other hand, many recent studies show that brain activity measured *during the choice process* is predictive of the magnitude of CIPC (Colosio et al, 2017; Jarcho, Berkman, & Lieberman, 2010; Kitayama et al, 2013; van Veen et al, 2009, Voigt et al, 2018). Unsurprisingly, key regions of the brain’s valuation and cognitive control systems are involved, including: the right inferior frontal gyrus, the ventral striatum, the anterior insula and the anterior cingulate cortex (ACC). Note that current neurocomputational theories of ACC suggest that it is involved in controlling how much mental effort should be allocated to a given task, based upon the derivation of the so-called “expected value of control” (Musslick et al., 2015; Shenhav et al., 2013). This is highly compatible with our results, under the assumption that pre-choice value reassessment is a controlled and effortful process that trades mental effort against choice confidence. We will pursue this computational scenario in subsequent publications.

Nevertheless, we consent that CIPC may be driven by both pre-choice value reassessment and post-choice cognitive dissonance reduction mechanisms. The quantitative contribution of the latter effect, however, may have been strongly overestimated. In our view, this is best demonstrated, though perhaps unintentionally, in the results of the “blind choice” study from Sharot and colleagues (Sharot et al., 2010). Here, participants rated items both before and after making a blind choice that could not be guided by pre-existing preferences (because the items were masked). Critically, although blind choice precludes any instrumental value refinement process, preferences were altered after the choice. Interestingly, the effect size is rather small, i.e. the ensuing CIPC magnitude was estimated to be around 0.07 ± 0.03. This is to be compared with the CIPC magnitude of the RCR and RRC conditions in two other studies by the same authors (Sharot et al., 2009, 2012), namely: 0.38 ± 0.08 (RCR condition, with non-blind choices) and 0.11 ± 0.06 (RRC condition, which was not included in the “blind choice” study). In other terms, CIPC under blind choice is smaller than the apparent CIPC that unfolds from the known statistical confounds of the free choice paradigm. Note that if this had not have been the case, then post-choice CIPC cognitive reduction effects would dominate and we would not have confirmed our predictions. Taken together, this series of studies clearly supports the notion that option re-evaluation occurs, at least partly, at the time of the decision.

## CONCLUSION

In conclusion, our results lend support to the hypothesis that choice-induced preference change is caused by an intra-choice refinement of option value representations that is motivated by difficult decisions, and undermine the established theory of post-choice cognitive dissonance resolution. We also demonstrate the relevance of meta-cognitive processes (cf. reports of choice confidence and certainty about value judgments) to choice-induced preference change. This contributes to moving forward the state of the 60-year-old research on the reciprocal influence between choice and subjective value.

## ETHICAL COMPLIANCE

Our analysis involved de-identified participant data and was approved by the ethics committee of the ICM (Paris, France). In accordance with the Helsinki declaration, all participants gave informed consent prior to commencing the experiment.

## DATA AVAILABILITY

All raw data will be made openly available upon request.

